# Cell-type specific projection patterns promote balanced activity in cortical microcircuits

**DOI:** 10.1101/2024.10.03.616539

**Authors:** Anno C. Kurth, Jasper Albers, Markus Diesmann, Sacha J. van Albada

**Author notes:** These authors contributed equally to this work (Anno C. Kurth, Jasper Albers).

## Abstract

The structure of neural networks provides the stage on which their activity unfolds. Models of cerebral cortex linking connectivity to dynamics have primarily relied on probabilistic estimates of connectivity derived from paired electrophysiological recordings or single-neuron morphologies obtained by light microscopy (LM) studies. Only recently have electron microscopy (EM) data sets been processed and made available for volumes of cortex on the cubic millimeter scale, exposing the actual connectivity of neurons. Here, we construct a population-based, layer-resolved connectivity map from EM data, taking into account the spatial scale of local cortical connectivity. We compare the obtained connectivity with a map based on an established LM data set. Simulating spiking neural networks constrained by the derived microcircuit architectures shows that both models allow for biologically plausible ongoing activity when synaptic currents caused by neurons outside the network model are specifically adjusted for every population. However, differentially varying the external current onto excitatory and inhibitory populations reveals that only the EM-based model robustly exhibits biologically plausible dynamics. Our work confirms the long-standing hypothesis that a preference of excitatory neurons for inhibitory targets, not present in the LM-based model, promotes balanced activity in cortical microcircuits.

## 1 Introduction

Microcircuits are the fundamental building blocks of the neocortex. Early observations identified a vertical compartmentalization into six layers [1] that, as a simplified concept, is still upheld to this day [2]. Horizontally, a modular organization consisting of repeating, stereotyped columns was suggested [3]. These concepts culminated in the notion of the “canonical microcircuit” that tries to reconcile the perceived generic structure of the neocortex with its function [4, 5]. The validity of this notion is subject to debate [6, 7] due to prominent non-uniformities in the composition of cortical tissue across species and areas within one species [8, 9]. Nonetheless, universal aspects of the area-specific microcircuits constrain cortical dynamics which ultimately underlie their function and thus are of fundamental importance for our understanding of the mammalian brain.

Single instances of local cortical circuits have been reconstructed experimentally [4, 10, 11, 12, 13], and their dynamics and information processing capabilities have been investigated theoretically [4, 14, 15, 16, 17, 18]. Their architecture is usually represented by a connectivity map specifying the probability for two neurons to establish a connection. These maps reduce the complicated circuitry to simple relations between cell types. Even though this approach neglects higher-order features like connectivity motifs [19, 20, 21], it enables investigations of the links between structural principles of local circuits and their dynamics. While downscaling the numbers of neurons and synapses distorts dynamical predictions [22], networks at realistic neuron and synapse counts provide a means to represent cortical dynamics including correlations faithfully.

Recent years have seen significant advances in the application of electron microscopy (EM) in neuroanatomy [23]. In particular, the adult networks for both sexes of *Caenorhabditis elegans* [24] and large volumes for *Drosophila melanogaster* (fruit fly) [25, 26] have been fully reconstructed. The work of Zheng et al. [25] subsequently formed the basis for obtaining for the first time a complete connectome of the adult fruit fly together with a comprehensive cell-type atlas [27, 28]. These efforts enabled further work on the statistical analysis of the connectivity [29] and a computational model of the whole fly brain at neuronal resolution [30]. In mammals, the wiring diagrams for a 0.0005 mm^3^ volume of mouse barrel cortex layer 4 [31] and for layer 2/3 pyramidal cells in a 0.003 mm^3^ volume of mouse primary visual cortex [32] were uncovered. Recently, Sievers et al. [13] obtained the full connectivity of all neurons in one barrel column of mouse barrel cortex containing about ten thousand neurons in a volume of roughly 0.2 mm^3^ of cortical tissue.

Combining advanced EM imagery with novel machine learning and data processing techniques recently enabled the reconstruction of local cortical networks in unprecedented detail at the cubic millimeter scale for mouse visual cortex [33] and human temporal cortex [34] handling petabytes of raw data. These reconstructions allow a more thorough look into the architecture of local cortical circuits beyond single columns than was previously possible. Due to technical limitations, the resulting connectomes are not complete: since axons require extensive manual proofreading, only a fraction of all synapses can be fully traced back to their source neurons. Consequently, a probabilistic description of connectivity is still needed.

In this study, we investigate the implications of newly obtained EM-based reconstructions for large-scale cortical modeling. To this end, we derive a layer-resolved, population-based connectivity map distinguishing excitatory and two inhibitory cell types from the subset of proofread neurons in a ∼ 1 mm^3^ EM reconstruction of mouse visual cortex [33]. As a reference, we compare with a model based on established light microscopy (LM) data by Binzegger et al. [11] which we construct according to identical principles. The choice of this dataset as a reference is motivated by its frequent use in computational modeling studies (see for example [35, 36, 15, 37, 38, 39]) and the long-lasting influence it enjoys to this day. Leveraging the explanatory power of the balanced random network paradigm [40, 41], we simulate spiking neural networks based on the two connectivity maps. Their comparison shows that, when assuming plausible synaptic interaction strengths, only the model based on the EM data exhibits biologically realistic, asynchronous irregular (AI) spontaneous activity while the LM-based model is prone to highly synchronized, biologically unrealistic states. Preliminary results have been presented in abstract form [42].

## 2 Results

To construct the microcircuit models, we first need to estimate the spatial range of local cortical connectivity between populations. This estimate is used to constrain the model sizes. Given these sizes, we determine probabilistic connectivity maps describing the local cortical circuitry. Finally, we perform spiking neural network simulations of the two models and link their dynamics back to their structure.

Binzegger et al. [11] estimate connectivity based on light-microscopic morphological reconstructions of 39 neurons from cat primary visual cortex, using a generalized version of Peters’ rule [43, 44]. Supplementing the data with previously obtained numbers of neurons and synapses [45, 46], the authors provide estimates for the numbers of synapses between neurons of various classes. For inhibitory neurons, the authors mainly focus on basket cells, leaving a certain fraction of inhibitory synapses unexplained.

The MICrONS data set [33] consists of calcium imaging of about 75 thousand neurons in the mouse visual cortex that is co-registered with an EM reconstruction of ∼ 1 mm^3^ including about 500 million synapses. The dataset has already been used in a number of scientific studies, for example to create a digital twin using an artificial neural network to predict neural responses and derive general wiring rules [47]. Another study employs unsupervised representation learning to uncover low-dimensional features of neural morphologies from the dataset [48]. For the present work, we exclusively focus on the EM reconstructions and only consider connections originating from 266 presynaptic neurons with fully reconstructed and proofread axons (see Section S1.1 for a precise description of the data used in this study). Also see Schneider-Mizell et al. [49] for a detailed analysis of a subset of the data with a view towards classical neuroanatomy distinguishing various inhibitory sub-classes revealing diverse and precise projection characteristics.

While the MICrONS data give direct access to *actual* neural connections, the data of Binzegger et al. [11] yield *potential* connectivity. In the following, we denote the model based on the MICrONS data set with ℳ_EM_, and the model based on the Binzegger data set with ℳ_LM_. In the two models, the cortical layers 2/3, 4, 5, and 6 are distinguished (henceforth, we refer to these as L2/3, L4, L5, and L6). Each layer contains one excitatory (E) and two inhibitory populations (Ib, Inb) corresponding to basket and non-basket cells. For the LM data set, we follow Izhikevich and Edelman [36], assigning unexplained inhibitory synapses to presynaptic non-basket cells. In the EM data set, all neurons can be straightforwardly mapped to one of the populations.

### 2.1 Number of synapses decays faster with distance than connection probability

We estimate the spatial organization of local cortical connections using both the distance-dependent mean number of synapses and the distance-dependent connection probability. Here, the connection probability is defined as the probability that two neurons establish at least one synapse. The connection probability consequently neglects the multiplicity of synapses between pairs of neurons.

We assume an exponential decay of the mean number of synapses *S*_*AB*_(*d*) between one presynaptic neuron of population *B* and one postsynaptic neuron of population *A* with horizontal somatic distance *d*:

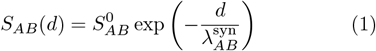

Here, 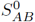 denotes the peak number of synapses, and 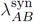 denotes the characteristic length of cortical connectivity between populations *B* and *A*. Taking the density *ω*(*d*) of distances *d* between pre- and postsynaptic neurons into account, the density of the expected mean number of synapses *s*_*AB*_(*d*) between a single neuron in presynaptic population *B* and all neurons in postsynaptic population *A* at distance *d* is *ω*(*d*) · *S*_*AB*_(*d*).

Likewise, we assume an exponential decay of the connection probability with distance *d* between individual neurons in the presynaptic population *B* and postsynaptic population *A*:

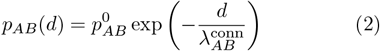

Here, 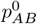 denotes the connection probability at zero distance. *p*_*AB*_(*d*) is the conditional probability for a neuron in population *B* to form at least one synapse with a particular neuron in population *A* given that the horizontal distance of their somata is *d* [51].

For ℳ_EM_, *S*_*AB*_(*d*) and *p*_*AB*_(*d*) can straightforwardly be extracted from the actual connectivity reported in the EM data. Figure 1a shows example fits of the density of the mean number of synapses, with resulting spatial decay constants 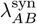 displayed in Figure 1A. We here assume a uniform distribution of neurons to approximate the true distribution of distances in the reconstructed volume, avoiding problems arising due to sparse data for some population pairs. (cf. Section 4.1). Further, we show example fits for the connection probability in Figure 1b and spatial decay constants 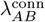 in Figure 1B. In agreement with the literature, we find that the spatial scales of local excitatory and inhibitory connectivity at the studied range are generally comparable, with a tendency for excitatory connection lengths to exceed the inhibitory ones [52, 53, 54, 20]. Furthermore, we observe that *S*_*AB*_(*d*) consistently exhibits smaller characteristic lengths than *p*_*AB*_(*d*). Calculating the fraction

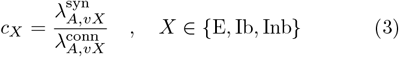

for all pairs of populations, where *v* is the layer of the presynaptic population and *X* is that population’s cell type, we observe a consistent average *c*^⋆^ = 0.7 (Figure 1C). Thereby, we can estimate 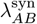 from 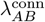, and vice versa.

**Figure 1:**
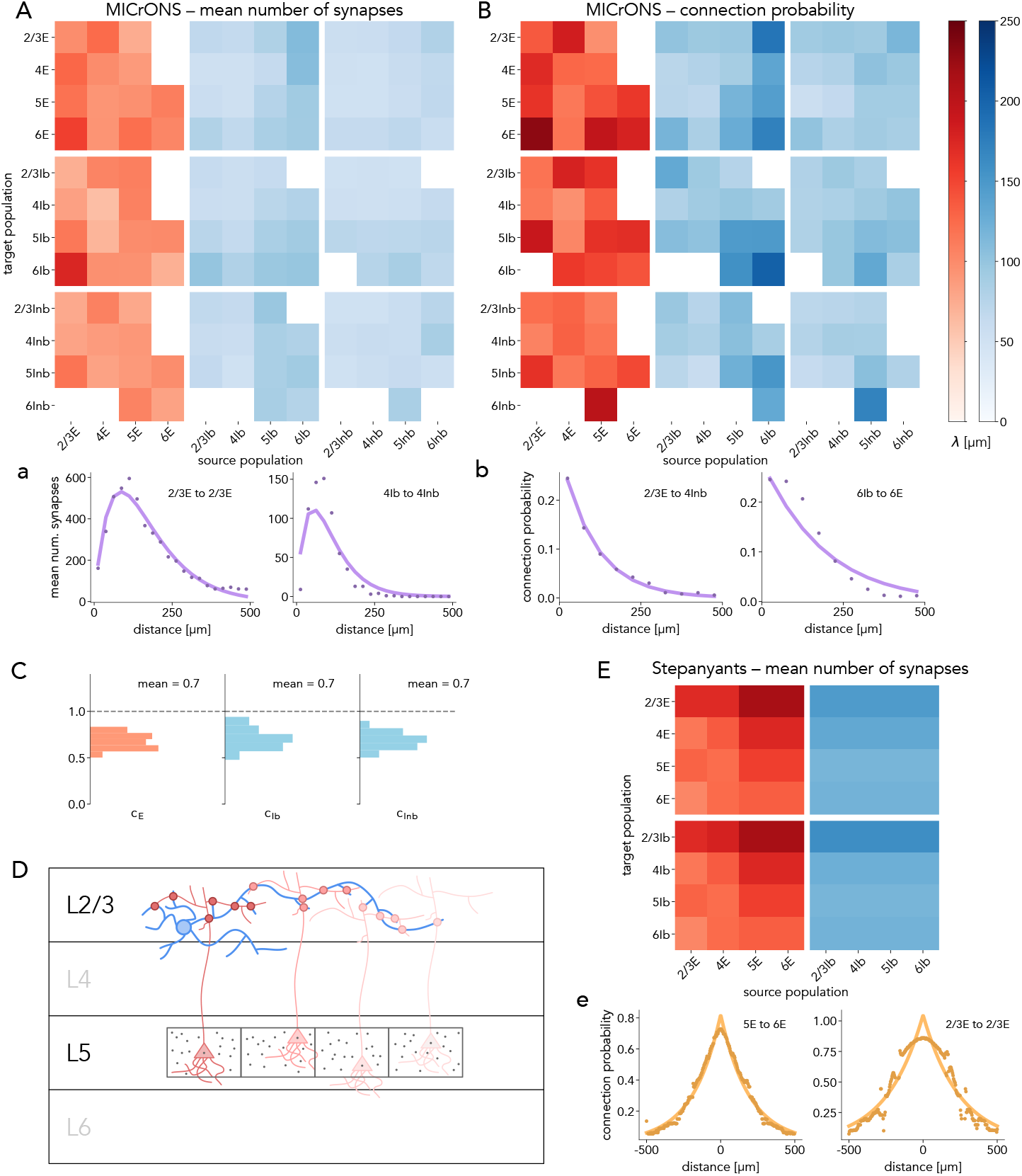
Spatial scale of actual and potential connectivity for the mean number of synapses and the connection probability. **A** Spatial scale of the mean number of synapses estimated from MICrONS data. Rows and columns organized according to population (E: excitatory, Ib: inhibitory basket, Inb: inhibitory non-basket) and cortical layer (2/3, 4, 5, 6). Colors in water tap notation: shades of red indicate excitatory, shades of blue inhibitory connections. White tiles indicate *<* 50 synapses. **B** Spatial scale of the connection probability estimated from EM data (actual connectivity). Same representation as in A. **C** Ratios between spatial scales of the mean number of synapses and the connection probability for actual connectivity from the EM data set. Histogram over all pairs of populations. **D** Illustration of calculation of potential connectivity from morphologies based on Peters’ rule. Location of presynaptic neuron (blue) of population *B* fixed. Postsynaptic neuron (red) of population *A* randomly placed in a box with side length Δ = 25 μm centered at horizontal position of presynaptic and original vertical coordinate of postsynaptic neuron. Position and rotation around vertical axis sampled *n* = 100 times (gray dots). Connection probability defined as *m/n* where *m* is the number of samples for which axon and dendrite are sufficiently close (red circles) at least once. **E** Spatial scale of the mean number of synapses estimated from data by Stepanyants et al. [50]. Excitatory-to-excitatory characteristic lengths estimated from connection probabilities and converted using *c*^⋆^. Remaining length scales estimated from expected number of potential synapses. Same color code as in A, B. **a**,**b**,**e** Example fits showcasing high-quality (left) and low-quality (right) curve fits (see Section S1.2 for overview of fits).

For ℳ_LM_, data on actual connectivity is not available. Instead, the spatial scale of the potential connectivity has been estimated solely based on the morphological reconstructions [50, 55]: a potential synapse between a pre- and postsynaptic neuron is registered if their axon and dendrite are sufficiently close (Figure 1D). With this approach, the authors estimate both the mean number of synapses and the connection probability. First, we assess the consistency of the LM potential connectivity with that from the EM data set. Using the provided morphologies, we reproduce the method (Figure 1D) to obtain an estimate of 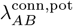 from potential connectivity. Comparing to the previously derived estimates from actual connectivity, 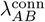, we find a good agreement across most pairs of populations (Figure S5).

Next, we calculate 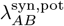 for all pairs of populations. In this methodology, we expect that the estimated number of potential synapses is prone to noise due to multiple adjacent segments of reconstructed neural processes being counted as locations of potential synapses. Thus, we use estimates of the connection probability for the pairings of populations where this information is available in [50], and then convert 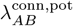 to 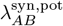 using *c*^⋆^. For all other pairs of populations, we directly estimate *S*_*AB*_(*d*). For pairs of populations with no data in [50], we make additional generalizing assumptions (Section S1.3). Example fits of this procedure are shown in Figure 1e, with the resulting 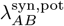 displayed in Figure 1E.

### 2.2 A common excitatory subcircuit of connectivity maps

Because our models ℳ_LM_ and ℳ_EM_ consider distance-independent mean connection probabilities *p*_*AB*_, the size of the represented cortical tissue and consequently the number of neurons and synapses must be fixed to derive the corresponding connectivity maps. To this end, we distribute neurons with realistic densities for each model in a template space. Assuming the distance-dependence of the mean number of synapses Equation 1, we determine the fraction of model-internal synapses for each pair of pre- and postsynaptic populations for circular patches of cortical tissue (Figure 2A). Here, model-internal synapses refer to incoming synapses originating from neurons in the model. The model radii 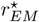 and 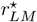 are then chosen such that the fractions of model-internal synapses averaged over pre- and postsynaptic populations are approximately 65 % (see Section 4.2), resulting in 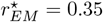 mm and 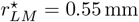.

**Figure 2:**
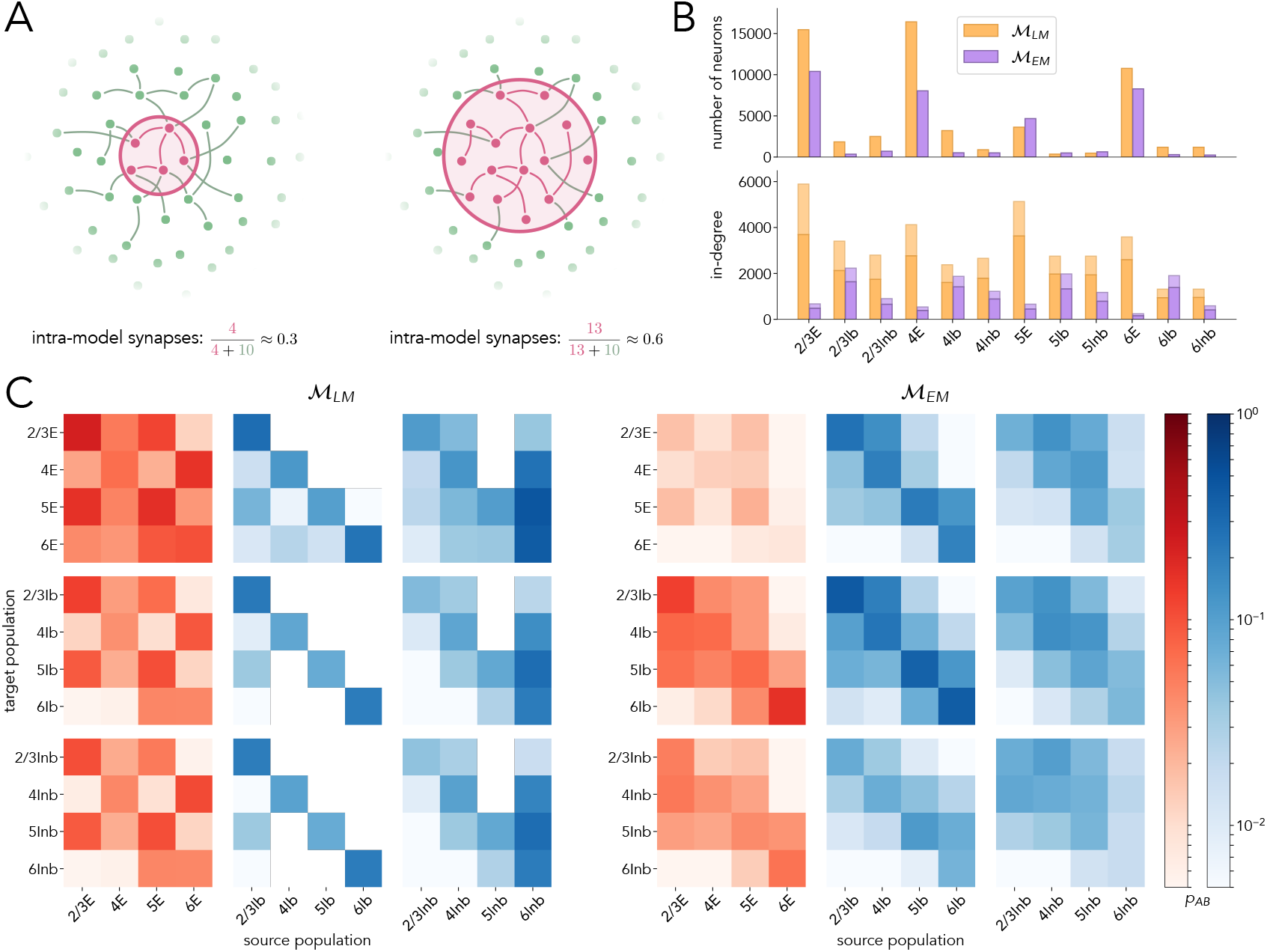
Model construction and connectivity maps. **A** Sketch illustrating the fraction of intra-model synapses depending on model size. **B** Numbers of neurons (top) and in-degrees (bottom) for both models with radii 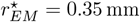 and 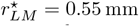. The model radii are chosen so that the fraction of model-internal synapses is approximately 65 % averaged over population pairs (Equation 5). For the in-degrees, solid bars indicate the number of intra-model synapses, while transparent bars (stacked) indicate synapses with presynaptic neurons not contained in the models. ℳ_EM_ consists of ∼ 37 thousand neurons and ∼24 million synapses; ℳ_LM_ consists of ∼ 58 thousand neurons and ∼ 244 million synapses. **C** Mean number of synapses between two arbitrary neurons of given source- and target populations (same representation as in Figure 1A).

We observe that ℳ_LM_ has a higher number of neurons in most populations, which is expected due to the larger area of cortical tissue covered by the model. Turning to connectivity, we observe that the populations of ℳ_LM_ have a significantly higher in-degree, i.e., number of incoming connections per neuron, except for 6Ib (Figure 2B). Consequently, the connection probabilities between populations are higher in ℳ_LM_ than in ℳ_EM_ (Figure 2C), suggesting a sparser recurrent connectivity in mouse visual cortex compared with cat. Still, common patterns can be identified: both models show an excitatory subcircuit between L2/3E and L5E, inhibitory basket cells project mainly within the same layer, and inhibitory non-basket cells preferentially target neurons in the same or higher layers. The main differences can be observed in the projection pattern of excitatory to inhibitory cells, as further discussed in Section 2.4.

### 2.3 ℳ_LM_ is highly synchronized while ℳ_LM_ exhibits AI activity

To investigate the dynamical properties of the constructed models ℳ_LM_ and ℳ_EM_, we instantiate and simulate spiking neural networks defined by the derived numbers of neurons and connectivity maps (see Section 4.3). Neurons are modeled as leaky integrate-and-fire units with conductance-based synapses using the neural simulation engine `NEST` [56]. Excitatory and inhibitory model neurons additionally include spike-frequency adaptation ([57], see Section 4.3). We constrain the recurrent synaptic weights using the data of Avermann et al. [58] for E, Ib, and Inb neurons of layer 2/3 in rat cortex. While this choice neglects layer- and species-specific diversity, it reduces model complexity and facilitates a direct comparison of the models. The weights are broadly consistent with a recently discussed hierarchy of connection strengths in mouse visual cortex [59] (see Section S1.4). Synaptic inputs from neurons not contained in the networks are modeled as excitatory conductance fluctuations in the form of Ornstein-Uhlenbeck processes with population-specific mean *μ*_*A*_, variance *σ*_*A*_, and time scale *τ*_*A*_ [60, 61]. The extrinsic drive to each neuron is statistically independent of all other drives. For a detailed model specification see Section S1.5.

The extrinsic drive to both network models is adjusted to approximately satisfy experimentally observed firing rates during spontaneous activity [62, 63] using the method suggested by Isbister et al. [61] (Table 7, Table 8). Concretely, *μ*_*A*_ and *σ*_*A*_ are chosen for each population *A* such that *σ*_*A*_ = *χμ*_*A*_ with cell-type independent *χ*. In the resultant state, the spiking activity of both networks exhibits biologically plausible characteristics: asynchronous irregular (AI) activity [64, 65] (Figure 3A,B), low firing rates obeying *ν*_Ib_ *> ν*_Inb_ *> ν*_E_ in most cases [66, 67, 68] (Figure 3C,F), broadly distributed coefficients of variation of the inter-spike intervals [69] (Figure 3D,G), and low synchrony assessed by co-fluctuations of neural membrane potentials [70] (Figure 3E,H). The dot displays only include neurons that spiked at least once in the observation interval, akin to experimental recordings where silent neurons remain unobserved.

**Figure 3:**
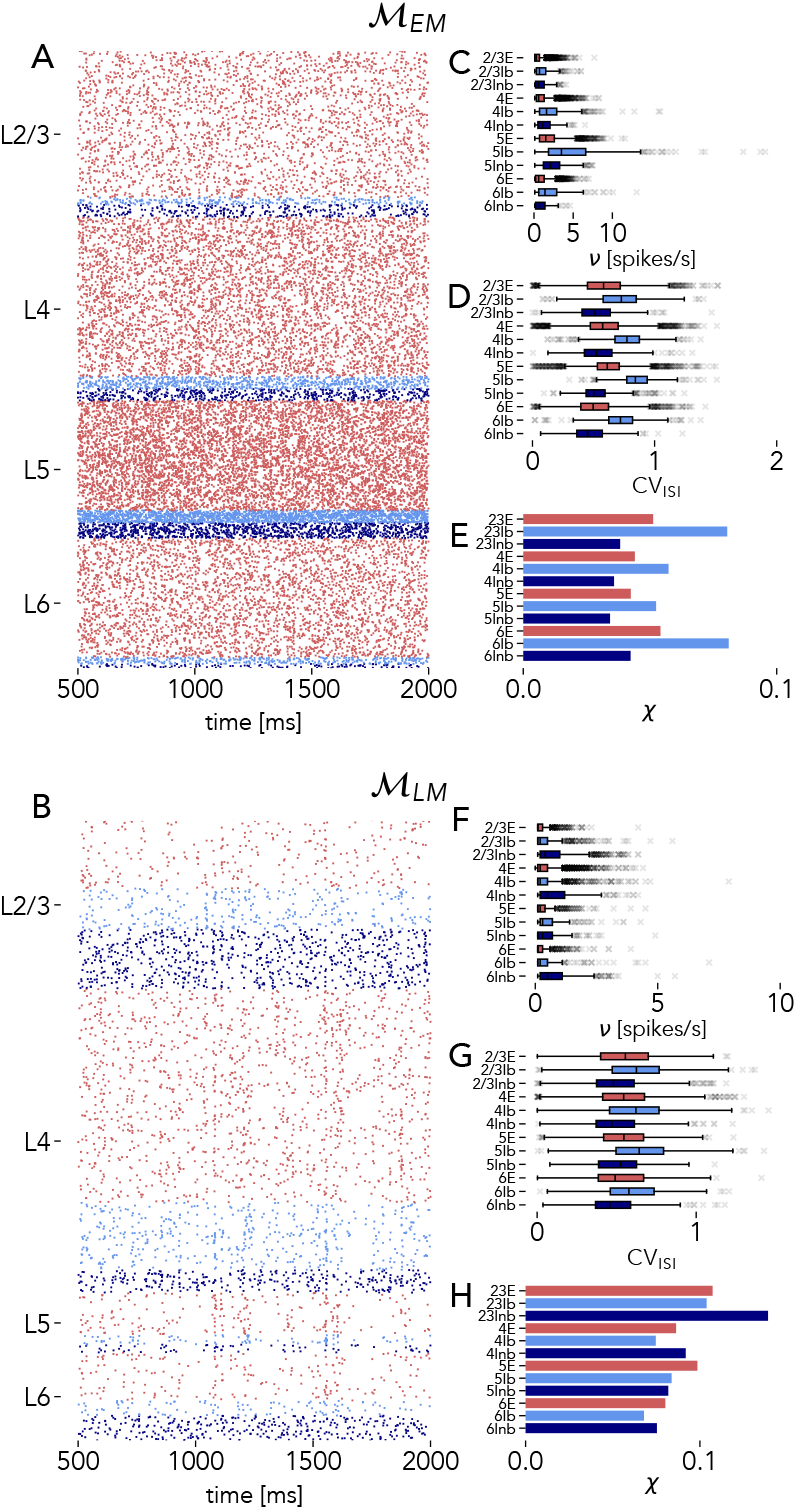
Model activity with optimized external input. Red indicates excitatory, light blue inhibitory basket, and dark blue inhibitory non-basket populations. The simulation gathers statistics over 10 s of biological time starting 0.5 s after simulation onset. **A, B** Raster plots (spikes marked as dots) of 50% of neurons that spiked at least once for the time stretch from 0.5 s to 2.0 s. **C–H** Population-resolved firing rates *ν*, coefficients of variation of the inter-spike interval CV_ISI_ (Equation 7), and synchrony *χ* (Equation 8).

To systematically investigate the dynamical repertoire of both models, we perform a parameter scan varying the mean drives *μ*_*E*_ and *μ*_*I*_ to the sets of excitatory and inhibitory populations. We find that ℳ_LM_ has only a narrow band of extrinsic drives for which the firing rates resemble experimentally observed activity: if *μ*_*I*_ is too large, the excitatory cells do not fire or have a vanishingly small firing rate, and if *μ*_*E*_ is too large, the network rapidly transitions to a highly synchronized state with high firing rates (Figure 4A-E left). In contrast, the activity of ℳ_EM_ smoothly depends on the extrinsic drive with plausible firing rates, irregular spiking, and low synchrony across a large domain of the parameter space (Figure 4A-E right). Additionally, ℳ_EM_ exhibits the paradoxical effect [71, 72, 73], where increasing *μ*_*I*_ decreases the firing rate of inhibitory basket cells. Note that also for ℳ_LM_ the inhibitory basket cells reduce their firing rate when increasing *μ*_*I*_ for large *μ*_*E*_ and small *μ*_*I*_. We do not consider this as a paradoxical effect, since for large *μ*_*E*_ and small *μ*_*I*_, ℳ_LM_ exhibits biologically implausible, highly synchronized activity, suggesting that in this case, the reduced inhibitory firing rate is due to a qualitative change in the dynamical state.

**Figure 4:**
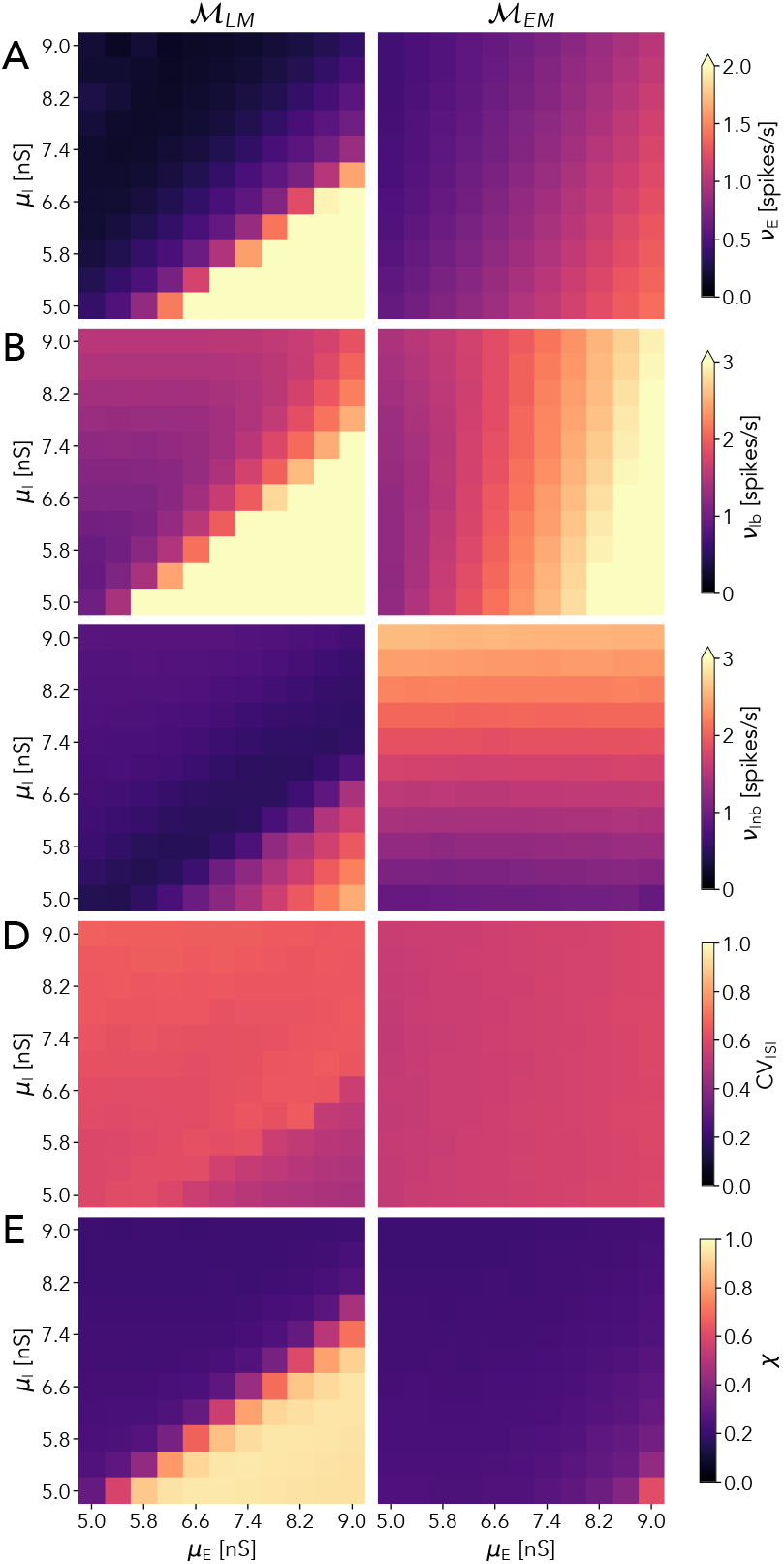
Model activity under variation of extrinsic drive. Simulation time of *T* = 10 s with a simulation period of *T*_pre_ = 0.5 s before data acquisition to avoid transients due to initial conditions. The mean input to all excitatory (*μ*_*E*_) and inhibitory (*μ*_*I*_) neurons is varied. Arrow at the top of color bars indicates that higher values are assigned the color of the maximum value. **A-C** Population-averaged firing rates of E, Ib, and Inb neurons. **D** Population-averaged coefficients of variation CV_ISI_ of the inter-spike interval (Equation 7). **E** Population-averaged synchrony *χ* (Equation 8).

### 2.4 Target specificity of excitatory neurons induces network-wide synchrony

What is the structural reason for this discrepancy between the models? Consider the specificity of projections from a given source population to excitatory versus inhibitory neurons in a target layer, which we term *target specificity* [15] (cf. Equation 9). The target specificity attains a positive (negative) value if a projection preferentially targets excitatory (inhibitory) neurons (Figure 5). For ℳ_LM_, all presynaptic populations have a positive target specificity. For ℳ_EM_, a more nuanced picture emerges where excitatory presynaptic populations preferentially target inhibitory neurons while there are inhibitory neurons of both cell types with positive as well as negative target specificities. This suggests that excitatory neurons that are more strongly innervated by recurrent connections, as is the case in ℳ_LM_, are the origin of pathological dynamics. We test this inℳ_EM_ by redistributing synapses from excitatory neurons that previously targeted inhibitory neurons such that they target excitatory neurons instead. This procedure conserves the total number of synapses. When a moderate number of synapses are redistributed, the target specificity of excitatory neurons becomes balanced (Figure 6A, left). The resulting model retains the biologically plausible dynamics across a large part of the parameter space (Figure 6B-F, left column). Increasing the number of synapses that are redistributed until the target specificity of excitatory neurons resembles that of ℳ_LM_ (Figure 6A, right), ℳ_EM_ shows qualitatively similar dynamics to ℳ_LM_ (Figure 6B-F, right column): for excitatory neurons, the transition between the firing regime at large *μ*_*E*_ and small *μ*_*I*_ and that at smaller *μ*_*E*_ and large *μ*_*I*_ becomes more abrupt. Moreover, the increased target specificity results in highly synchronous activity for a larger portion of the parameter space. In addition, Note, the CV_ISI_ increases in the part of the parameter space where highly-synchronized activity can be observed. Thus, the biologically implausible activity of ℳ_LM_ can in part be explained by the underlying positive target specificity, which in turn stems from the assumptions of the reconstruction of the local cortical circuit.

**Figure 5:**
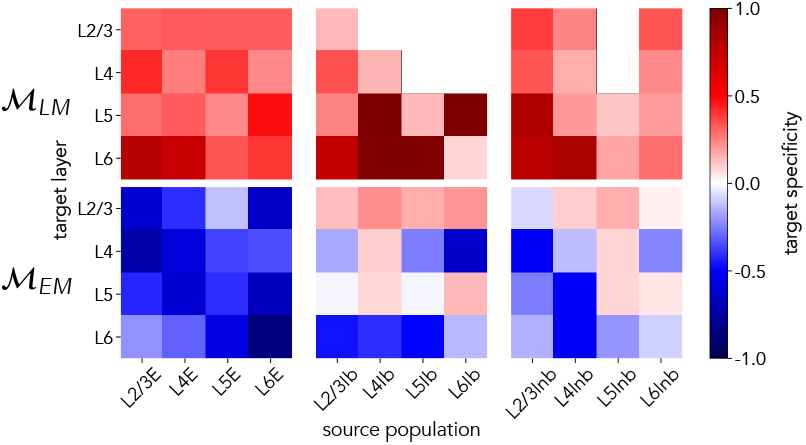
Target specificity for both network models. Positive values indicate a preferential targeting of excitatory neurons in a layer by a given source population; negative values indicate preference for inhibitory targets.

**Figure 6:**
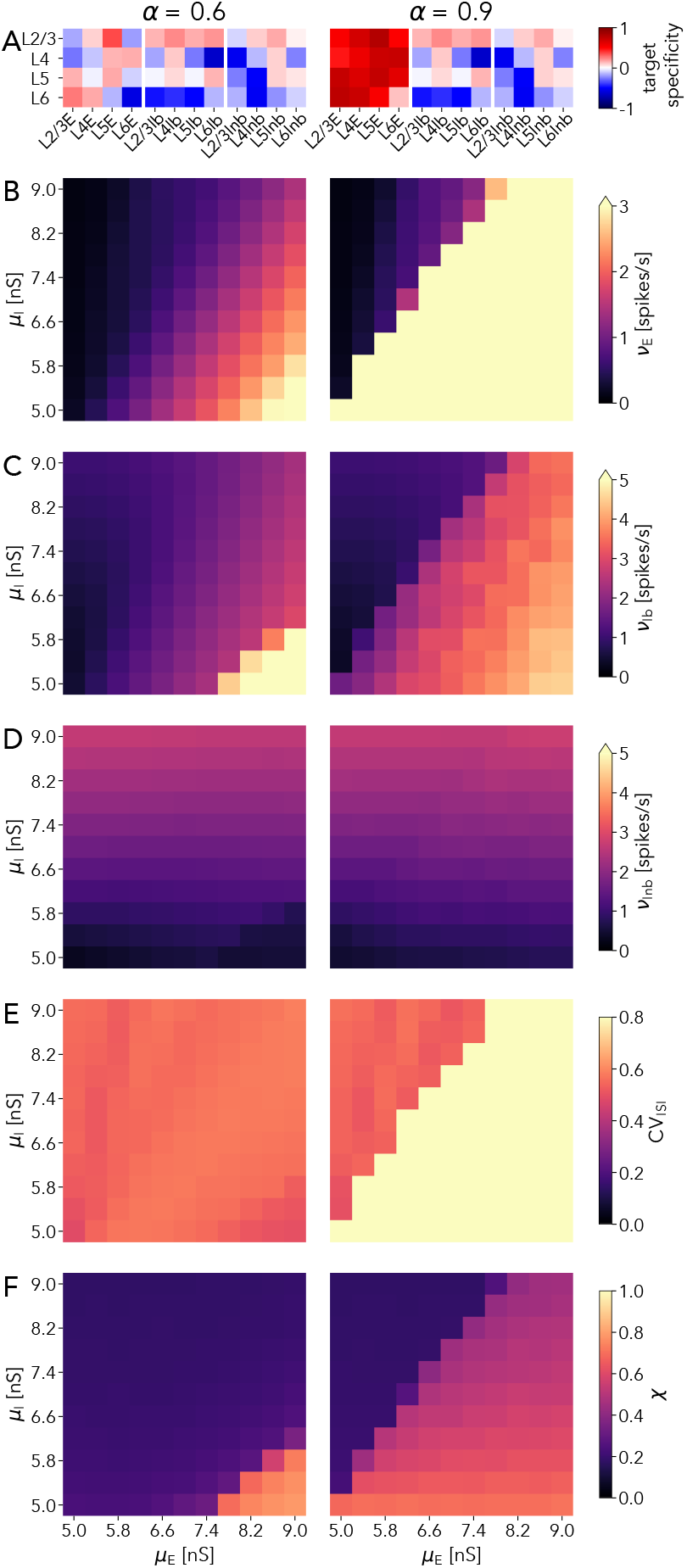
Effect of increased target specificity in ℳ_EM_. Parameter *α* ∈ [0, 1] linearly controls redistribution of synapses onto excitatory neurons leading to an increase in target specificity (cf. Section 4.5). *α* = 0 corresponds to the original model, *α* = 1 implies that all synapses target excitatory neurons. Color code in top panels as in Figure 5, in bottom panels as in Figure 4.

## 3 Discussion

In the present work, we revisit local cortical microcircuit architectures. Based on two reconstruction techniques—light microscopy (LM) and electron microscopy (EM)—we compare cortical connectivity and study implications for cortical dynamics using spiking neural network models. Our results show that strong recurrent innervation of excitatory neurons leads to pathological dynamics in the LM-based model. Consequently, the LM-based connectivity needs to be modified to allow for biologically plausible activity over a wide range of input parameters. Conversely, the EM-based model exhibits balanced, biologically plausible activity for a wide range of extrinsic drives without the need for further modifications. This can be understood in terms of the target specificity: excitatory neurons preferentially target inhibitory ones, while inhibitory neurons have a diverse targeting pattern.

Such layer-specific projection patterns are in agreement with the study by Potjans and Diesmann [15] who amended connectivity derived from Binzegger et al. [11] with physiological connectivity data from several species and cortical areas, motivated by a discrepancy in target specificity between data sources (see their Figures 4 and S3). This reorganization leads to a stronger targeting of inhibitory neurons in certain layers by excitatory neurons, which is essential for asynchronous and irregular activity with realistic firing rates. Our results confirm the hypothesized negative target specificity of especially excitatory projections using reconstructions from a single brain.

Neither ℳ_EM_ nor ℳ_LM_ should be understood as detailed models of the mouse or cat visual cortex, respectively. For this, more fine-grained anatomical and physiological data regarding, for example, orientation specific preferential targeting or the modular lateral organization should be integrated (see for example Antólik et al. [39] and Billeh et al. [18]). The models presented here should rather be understood as scaffold microcircuits that highlight general dynamical properties of the respective network.

### 3.1 Modeling paradigm

The approach to modeling neural circuits employed here rests on the seminal insights from balanced random networks [40, 41, 74]. These networks have since been extended to multiple populations taking into account biological details such as the cell-type specific organization of cortex [15, 75], multiple inhibitory populations [75, 76, 38], or morphologically detailed neuron models [16, 61] to study cortical dynamics and function. Here we present a random network model constrained by EM-based connectivity data of mammalian cortex. While derived from a specific connectome, the obtained probabilistic connectivity rules represent the general connectivity of a template mouse visual cortex and allow for efficient instantiations of neural network models in distributed computers [77]. As such, they are amendable by future EM reconstructions and provide a starting point for generic cortex modeling. Using the derived characteristic spatial extents of connectivity, the model can be promoted to a spatially organized network that is scalable to sizes beyond the local cortical circuit. Such networks exhibit rich dynamics across space [78, 79, 80] and enable investigations of, for example, activity and function in visual cortical areas [17, 18, 81].

Alternatively, bio-realistic modeling can be based on the specific, completely reconstructed circuitry of one animal. In terms of mammalian local circuits spanning all layers, these data are only available for one column of mouse barrel cortex [13]. Advanced efforts combine physiological measurements of the activity of a piece of tissue with a subsequent EM reconstruction, allowing for self-consistent investigations of the relationship between the structure, dynamics, and function in a particular brain (as for example done for the MICrONS dataset [33], see also Arkhipov et al. [82] for an extended discussion). In contrast, the abstractions employed in the probabilistic approach put forward here expose principles of cortical architecture and processing.

### 3.2 Spatial decay of connection probability and mean number of synapses

Different spatial scales of local cortical connectivity in mouse visual cortex and cat V1 (Figure 1A, E) are not surprising given interspecies differences. However, also the methodologies used to determine the decay of connection probability with distance differ between the data sets. While LM data only allow estimating the potential connectivity, the EM data allow deriving the spatial extent of actual connectivity based on established synapses and accurate neuron positions as well as potential connectivity from the provided morphologies. Comparing the two methodologies for deriving the distance-dependent connection probabilities from the EM data set (Figure S5), we find a general agreement between the estimates of characteristic length scales.

Further, using the detailed information provided by the EM data, we compare the spatial decay of connection probability to that of the mean number of synapses. For both quantities, we assume an exponential decay with distance, which provides accurate estimates for a wide range of pre- and postsynaptic pairs of populations (Figure S1, Figure S2). We show that the mean number of synapses exhibits a decay with a consistently shorter characteristic length for all considered pairs of populations (Figure 1C). The reason for this difference may be that synapses between a given pair of neurons ‘beget’ more synapses, leading to long-tailed distributions of synaptic multiplicity [83]. Numbers of neuron pairs with high synaptic multiplicity would then fall off faster with distance than numbers of neuron pairs with low synaptic multiplicity, causing the observed difference in spatial decay between connection probability and the mean number of synapses.

The exponential model for the connection probability with distance (Equation 2) is in agreement with the experimental data as well (Figure S3). For distances above about 100 μm, Markov et al. [84] suggest an exponential decay with *λ* = 230 μm for the fraction of labeled neurons from retrograde tracing in macaque visual cortex. This finding is consistent with our results, whether fractions of labeled neurons are more reflective of numbers of synapses or of connection probabilities, since both follow an exponential decay with distance. Similarly, Boucsein et al. [51] show that also distance-dependent connection probabilities in rat somatosensory cortex obtained by physiological means follow an exponential decay with *λ* = 330 μm. Thus, our study qualitatively confirms previous work on the decay of connection probability between individual neurons with distance.

### 3.3 Local connectivity of mouse visual cortex is less dense

While both models derived here have a comparable neuron density per mm^2^ surface of cortex (with the density of ℳ_EM_ exceeding the density of ℳ_LM_), our analysis suggests a stark contrast for the density of local synapses. After correcting for the omission of extra-model synapses with cell body in the visual cortex, our analysis yields that around ∼ 85% of synapses in cat V1 stem from V1-internal projections (total number of synapses in cat V1 taken from [46]). This is consistent with the fraction of internally labeled neurons in retrograde tracing experiments reported by Markov et al. [84] for macaque V1 (a species with a brain surface area of the same order of magnitude).

However, a similar calculation for mouse visual cortex based on the MICrONS data suggests that only ∼ 20% of synapses are internal to that area (total number of synapses taken from the MICrONS data set, consistent with figures reported in [85] for mouse visual cortex). This is supported by previous reports highlighting sparsity of recurrent connections estimated by physiological means in single layers of mouse visual cortex [86] or throughout the cortical column [87] within a radius of 100 μm from the patched post-synaptic cell. These probabilities, combined with the exponential decay of connection probabilities with distance described here, make the comparatively small number of recurrent synapses internal to mouse visual cortex appear reasonable. Additionally, the large difference between mouse and cat is plausible in light of the extremely high inter-area connectivity targeting the mouse visual cortex reported by Gămănuţ et al. [88]. A comparative analysis by Magrou et al. [89] revealed that, while in species with larger brains like the marmoset and the macaque, V1 primarily receives inputs from higher visual areas, mouse visual cortex is targeted by all cortical areas investigated so far (including visual, other sensory, and motor areas). A recent analysis of large-scale neural recordings in mouse cortex indicates that these projections are sufficiently strong to modulate the activity in visual cortex and imprint behavioral variables beyond visual stimuli [90]. Taken together, this is in agreement with the observed sparse recurrent connectivity in ℳ _EM_.

### 3.4 Target specificity controls dynamical repertoire

Instantiating and simulating spiking neural network models based on the derived connectivity maps and spatial scales, we find that both the EM- and LM-based models can exhibit biologically plausible activity (Figure 3). However, systematically varying the background drive to excitatory and inhibitory populations reveals that ℳ_LM_ only behaves in a biologically plausible way for a narrow band of the parameter space: the model is prone to completely silent excitatory populations or network-wide strong, pathological oscillations. In contrast, ℳ_EM_ is well-behaved across a wider range of drive parameters, a necessary prerequisite for robust cortical computation. Furthermore, ℳ_EM_ reproduces the experimentally observed paradoxical effect for inhibitory basket cells [72].

The differences in dynamical behavior between the two models can be explained by cell-type-specific targeting patterns. Our results indicate that a strong preference of excitatory populations for targeting excitatory populations leads activity that is prone to instabilities: for ℳ_LM_, the aforementioned targeting pattern is observed (Figure 5); redistributing synapses in ℳ_EM_ to match the targeting pattern of ℳ _LM_ leads to qualitatively similar dynamics (Figure 6).

The preferential targeting of inhibitory neurons by excitatory neurons observed in ℳ_EM_ thus stabilizes the activity in the presence of strong external drive. Johnson and Burkhalter [91] showed that inter-area projections in rodents are predominantly excitatory and overwhelmingly target excitatory neurons. Other work highlights that in cortical feedback pathways in the mouse, inhibition is less effectively recruited than in feedforward pathways, judged from excitatory postsynaptic currents in parvalbumin-expressing interneurons compared with those in pyramidal cells [92]. The stabilizing effect of the recurrent connectivity might, in the behaving mouse, thus be required for the integration of inputs from a high number of brain areas in the lower visual cortical areas.

The notion of target specificity put forward here paints with a broad brush, highlighting population-level connectivity schemes governing population dynamics. The analysis of Schneider-Mizell et al. [49], distinguishing various sub-classes of inhibitory neurons, studies target specificity on a finer level, revealing highly specific projection patterns. Future work could aim to reconcile our approach with the specialized targeting patterns uncovered by Schneider-Mizell et al. [49].

### 3.5 Conclusion

In this work, we provide openly accessible connectivity maps and models of the cortical microcircuit based on established LM and recent EM reconstructions. As such, the obtained maps and models provide platforms for future modeling studies of local cortical networks. Possible extensions include taking into account more detailed single-neuron parameters and connection weights, the distribution of the multiplicity of synapses between pairs of pre- and one post-synaptic neurons, higher-order connectivity motifs, distance-dependent connectivity, functionally specific external input, neuronal morphologies, and functional properties such as plasticity.

Both models, ℳ_LM_ and ℳ_EM_, reduce the intricate morphology and complex response patterns of nerve cells to leaky integrate-and-fire point neuron models with conductance-based synapses and, in the case of excitatory cells and inhibitory non-basket cells, two adaptation currents. Given this level of description, the EM-based model outperforms the LM-based model in terms of biological plausibility of the emerging activity due to differences in cell-type-specific targeting patterns. This suggests that the EM-based connectivity is to be preferred as a starting point in future modeling of local cortical circuits. Our study confirms by the analysis of direct anatomical measurements the prediction of the existence of a negative target specificity of excitatory projections realized in the microcircuit model of Potjans and Diesmann [15].

## 4 Methods

### 4.1 Spatial connectivity

To obtain estimates for the characteristic lengths of local cortical connectivity, we assume an exponential decay for the mean number of synapses with lateral distance between somata (Equation 1). The EM reconstruction allows for determining the density of the mean number of actual synapses *s*_*AB*_(*d*) between a single presynaptic proofread neuron and all postsynaptic neurons at inter-somatic distance *d* for pairs of the considered cell types. Using the distribution of uniformly sampled points in a rectangle with distance *d* [93], we approximate the number of potential postsynaptic partners *ω*(*d*) given the dimensions of the reconstructed volume. We then fit

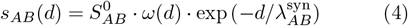

to obtain *λ*_*AB*_ between presynaptic population *B* and postsynaptic population *A*.

Additionally, we estimate the distance-dependent connection probability, i.e., the probability of establishing at least one synapse. We define the connection probability *p*_*AB*_(*d*_1_, *d*_2_) between a presynaptic neuron of population *B* and a postsynaptic neuron of population *A* that have a horizontal inter-somatic distance *d* with *d*_1_ ≤ *d* ≤ *d*_2_ as the number of connected neuron pairs divided by the total number of neuron pairs in the same range of distances. Here, the total number of neuron pairs is given by the product of the number of neurons of populations *A* and *B* within the range of distances. We also assume an exponential decay for the continuous *p*_*AB*_(*d*), albeit with a different characteristic length scale (Equation 2). The quantity 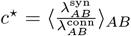 measures the discrepancy between the distance dependence of the mean number of synapses and the connection probability based on the EM data set.

For the LM data set, Stepanyants et al. [50] derive the connection probability using potential connectivity, defined as the probability that a source and target neuron of given cell types at inter-somatic distance *d* form at least one synapse based on morphological reconstructions of single neurons. This connection probability is given for a subset of pairs of populations. For these, we again fit Equation 2, which we convert to an estimate of the characteristic length of the mean number of synapses 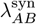 using *c*^⋆^. For a different subset of pairs of populations, Stepanyants et al. [50] derive the distance-dependent mean number of synapses *S*_*AB*_(*d*) under further assumptions, which we fit using Equation 1 to also obtain *λ*^syn^. These fits are displayed in Figure S4. Finally, for pairs of populations in our model where Stepanyants et al. [50] provide neither of the two quantities, we generalize from the existing estimates as detailed in Section S1.3.

Further, we assess the consistency of the LM potential connectivity with the potential and actual connectivity from the EM data. For this, we use the same morphology-based ansatz to derive a connection probability based on potential connectivity from the EM data. Compared to [50], we use more single-cell reconstructions (266 compared to 41), and fewer cell positions within each box (100 compared to 1000). Then, we compare the resulting length scales of spatial connectivity with estimates from the actual connectivity derived above. For the EM data set, the two methodologies result in estimates of 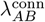 that are comparable across all pairs of populations.

In all cases, `scipy.optimize.curve fit` [94] performs non-linear least-squares fits using the Levenberg-Marquardt algorithm [95].

### 4.2 Connectivity maps

To constrain the size of the models, we calculate

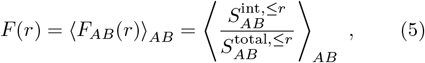

the fraction of intra-model synapses for a circular patch of cortex with radius *r*. To determine the expected number of synapses from population *B* to population *A* of neurons within a circular patch with radius *r*, 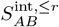, we uniformly distribute neurons with realistic densities in the patch and calculate the number of synapses according to Equation 1. Similarly, the total number of synapses between the populations, 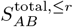, is determined by increasing the sampling radius of the presynaptic population to 3*r*. The model radius *r*^⋆^ for both models is determined so that *F* (*r*^⋆^) ≈ 65%.

We calculate the number of synapses between populations *A* and *B* internal to the model as

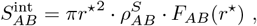

where 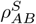 is the density of synapses from population *B* to population *A* extracted from the respective data set, and finally the distance-independent connection probability between a neuron in presynaptic population *B* and a neuron in postsynaptic population *A* as

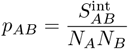

where *N*_*A*_, *N*_*B*_ are the number of neurons in populations *A* and *B*, respectively.

Population pairs having insufficient data to constrain the characteristic length scales (cf. S2) are neglected in determining the model sizes, but they are nevertheless connected according to the (scaled) total number of synapses in the respective data sets. Thus, also populations without a fitted length scale are connected, unless the corresponding number of reported synapses is zero.

### 4.3 Network model and single-neuron dynamics

Using the derived connectivity maps we instantiate network graphs for the two models using a pairwise Bernoulli connection rule with source- and target-population-specific connection probabilities *p*_*AB*_ (see the user-level documentation of NEST for an ontology based on [96]). The resulting networks are directed versions of Erdő s-Rényi graphs [97].

Single neurons are modeled as leaky integrate-and-fire units with conductance-based synapses and additional conductances that implement spike-frequency adaptation (a). The membrane potential of all units evolves as

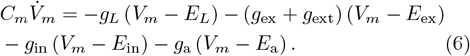

with the membrane capacitance *C*_*m*_, leak conductance *g*_*L*_ and reversal potentials *E*_*L*_, *E*_ex_, *E*_in_. The conductances obey

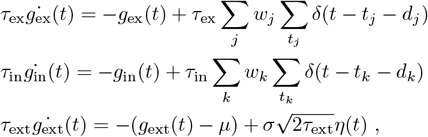

where *j* (*k*) is a presynaptic neuron, *t*_*j*_ (*t*_*k*_) its spike times, *w*_*j*_ (*w*_*k*_) the weight on a target neuron, *d*_*j*_ (*d*_*k*_) is the connection delay, *τ*_ex_, *τ*_in_ are the synaptic time constants, *μ* is the mean and *σ* the standard deviation of the extrinsic drive modeled as an Ornstein-Uhlenbeck process (*η* denotes white noise), and *τ*_ext_ its timescale. The timescales of response conductances and the extrinsic conductance modulation can differ since the latter entails slower fluctuations from the not explicitly modeled neural tissue. For all simulations, we fix *σ/μ* = *χ* = 0.2 [60] while the mean varies between populations and experiments. If *V*_*m*_ crosses the threshold *V*_th_ at time *t*^*′*^, the unit emits a spike with this time-stamp, is reset to *V*_reset_, and is kept at this value for *τ*_ref_. The conductances of the spike-frequency adaptation and relative refractoriness evolve according to

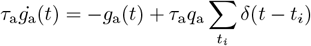

where *t*_*i*_ are the timings of the spikes emitted by the neuron, *τ*_a_ the timescale of the adaptation dynamics, and *q*_a_ the change in adaptation or relative refractoriness upon spike emission.

The neuron models used in this study are specified using `NESTML` [98]. The state of the neurons (Equation 6) is integrated with an embedded Runge-Kutta-Fehlberg 4(5) method [99]. The Langevin equation describing the extrinsic drive is integrated using an exact scheme [100, 101].

### 4.4 Analysis of dynamical data

The firing rates are determined for each neuron by counting the spikes and dividing by the observation time. Coefficients of variation of the inter-spike intervals (ISIs) for each neuron are given by

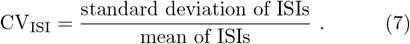

Both quantities are calculated using the Electrophysiology Analysis Toolkit (`elephant`[102]). To determine the synchrony *χ* of each population, we record the membrane potential of 50 neurons per population and calculate

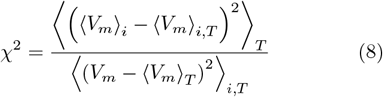

where ⟨…⟩_*i*_ denotes the average over the neurons in one population, and ⟨…⟩_*T*_ denotes the time average.

### 4.5 Target specificity

For a presynaptic population *B* and a layer *v*, the target specificity is given by

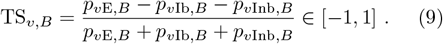

A positive (negative) value indicates that *B* preferentially targets excitatory (inhibitory) neurons in layer *v*.

We redistribute synapses between populations to increase the value of the target specificity using a control parameter *α* ∈ [0, 1] while keeping the total number of synapses fixed. On the level of the mean number of synapses, this amounts to:

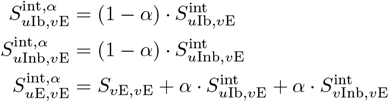

This definition implies that *α* = 0 leaves the connectivity maps (and thus the target specificity values) unchanged, while for *α* = 1 all synapses exclusively target excitatory populations (TS = 1).

## Code and data availability

The data and code to reproduce the results are openly available on Zenodo (https://doi.org/10.5281/zenodo.13886620).

## Author Contribution

Conceptualization: AK, JA Methodology: AK, JA, SvA, MD; Software: JA, AK; Formal analysis: JA, AK; Writing - original draft: AK, JA; Writing - review & editing: AK, JA, SvA, MD; Visualization: JA, AK; Supervision: SvA, MD; Funding acquisition: SvA, MD

## Acknowledgments

This project has received funding from NeuroSys as part of the initiative “Clusters4Future” by the Federal Ministry of Education and Research BMBF (03ZU1106CB), the European Union’s Horizon Europe Programme under the Specific Grant Agreement No. 101147319 (EBRAINS Project) and the Priority Program (SPP 2041 “Computational Connectomics”) of the Deutsche Forschungsgemeinschaft. The authors gratefully acknowledge the computing time granted by the JARA Vergabegremium and provided on the JARA Partition part of the super-computer JURECA at Forschungszentrum Jülich (computation grant JINB33).

## S1 Supplementary materials

### S1.1 Electron Microscopy data

We use the electron microscopy reconstructions from the MICrONS data set[1]. More precisely, we use the dataset `minnie65_phase3_v1` with timestamp `version=658`. As query tables, we use `allen_v1_column_types_slanted`, `baylor_gnn_cell_type_fine_model_v2`, `aibs_soma_nuc_metamodel_preds_v117`. Finally, we only consider presynaptic neurons with manually proofread axons, employing the query `proofreading_status_public_release` where we set the status `status_axon==‘extended’`.

### S1.2 Population-specific characteristic lengths

We fit an exponential decay of the mean number of synapses using Equation 1 and the connection probability using Equation 2 to the actual and potential connectivity derived from EM data to obtain the population-specific characteristic lengths *λ*_*AB*_. Figure S5 compares the length scale of the connection probability obtained from actual and potential connectivity. We find that the two methods produce similar estimates:

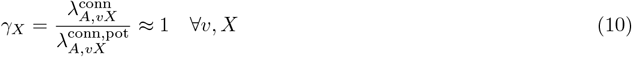

where *v* is the layer and *X* is the cell type of the presynaptic population.

**Figure S1:**
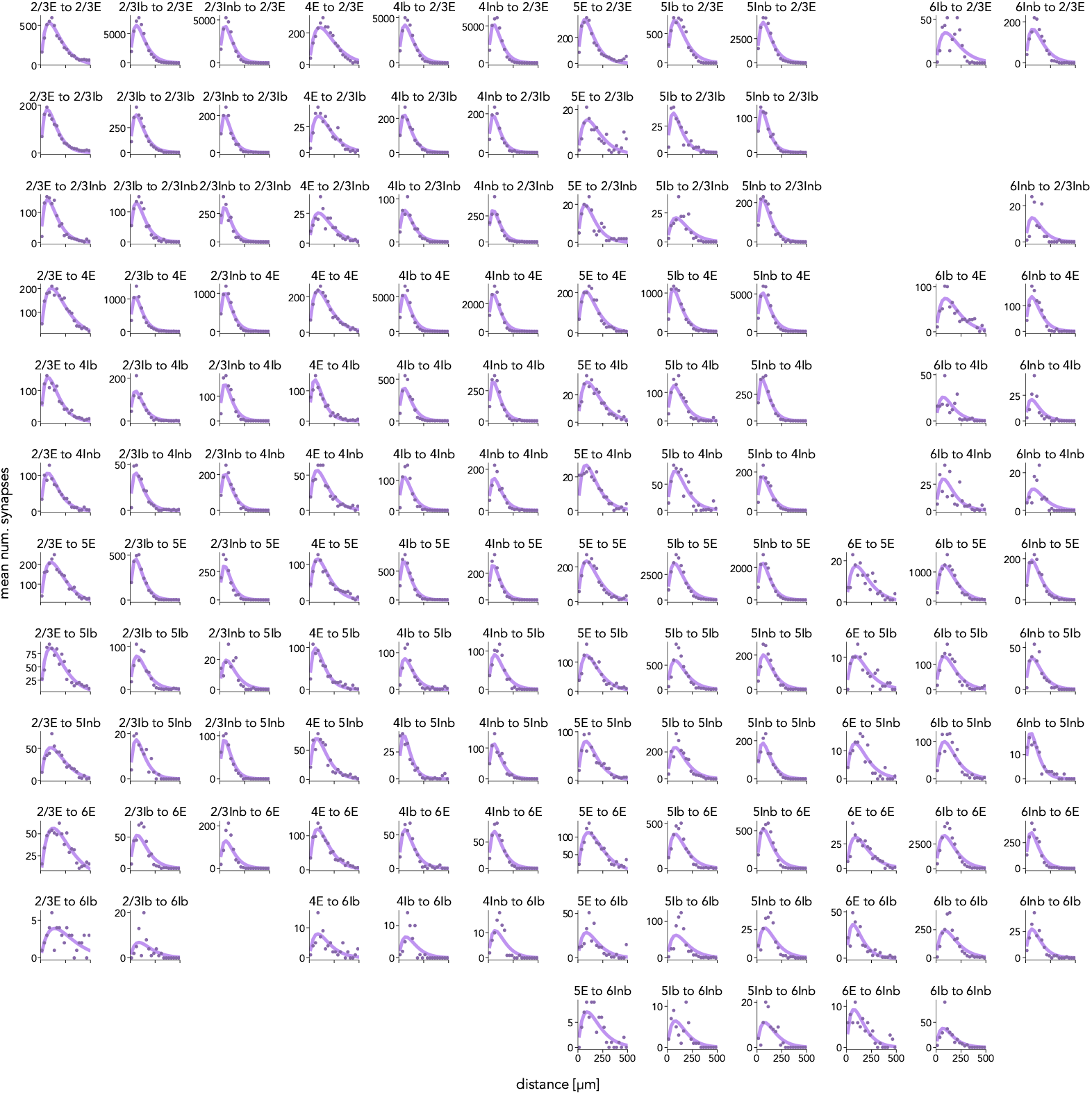
Density of mean number of synapses between pre- and postsynaptic populations from EM data, actual connectivity. Purple dots indicate experimentally observed number of connections at distance *d*, purple curves are fits to the expected number of connections given the density of the mean number of synapses of one presynaptic neuron with all possible partners from the postsynaptic population at distance *d* (Equation 4). No fits for combinations of populations where the observed number of synapses is smaller than 50.

**Figure S2:**
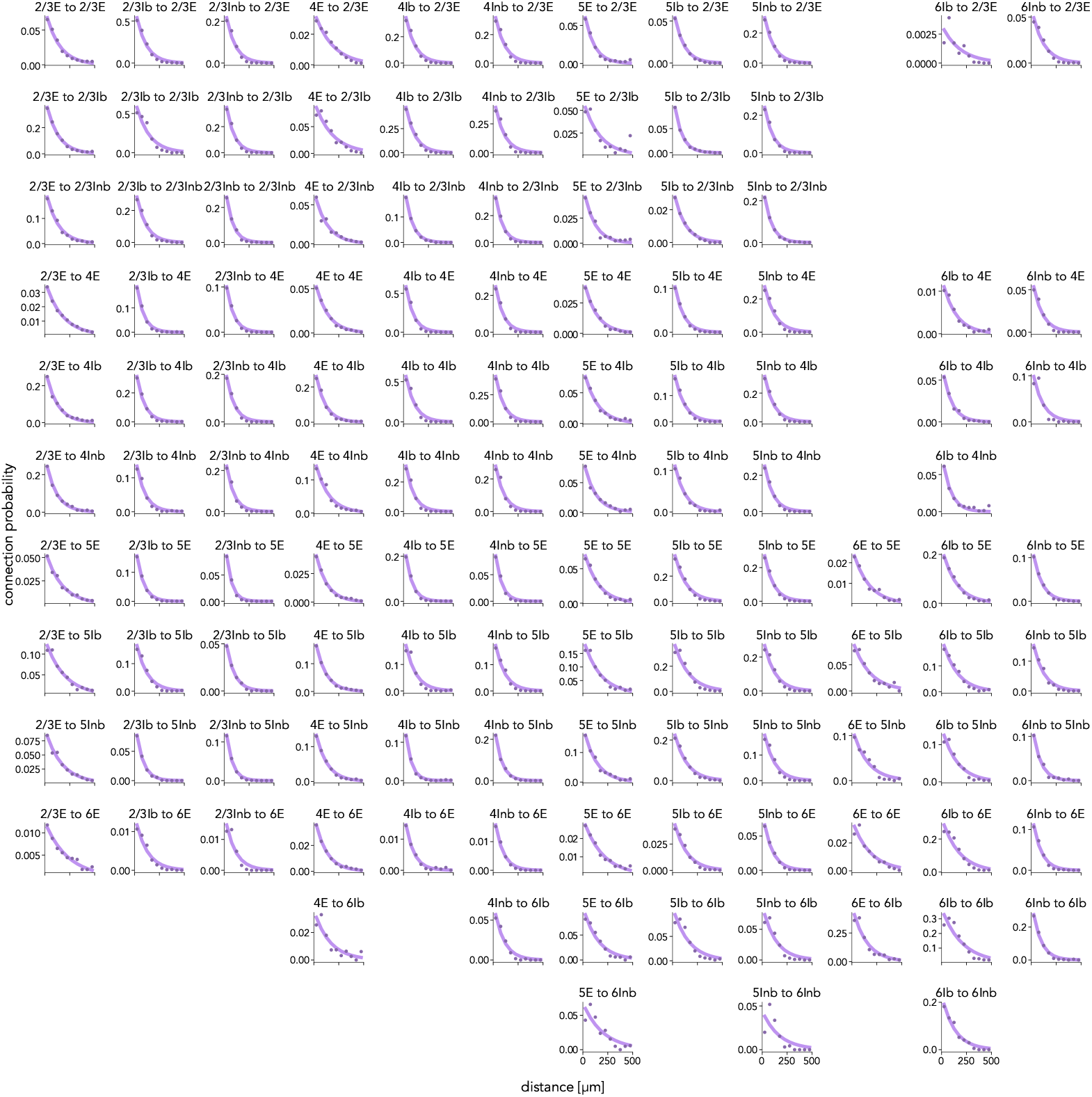
Connection probability between pre- and postsynaptic populations from EM data, actual connectivity. Purple dots indicate experimentally observed connection probability at distance *d*, purple curves are fits to the expected connection probability (Equation 2). No fits for combinations of populations where the observed number of connections is smaller than 50.

**Figure S3:**
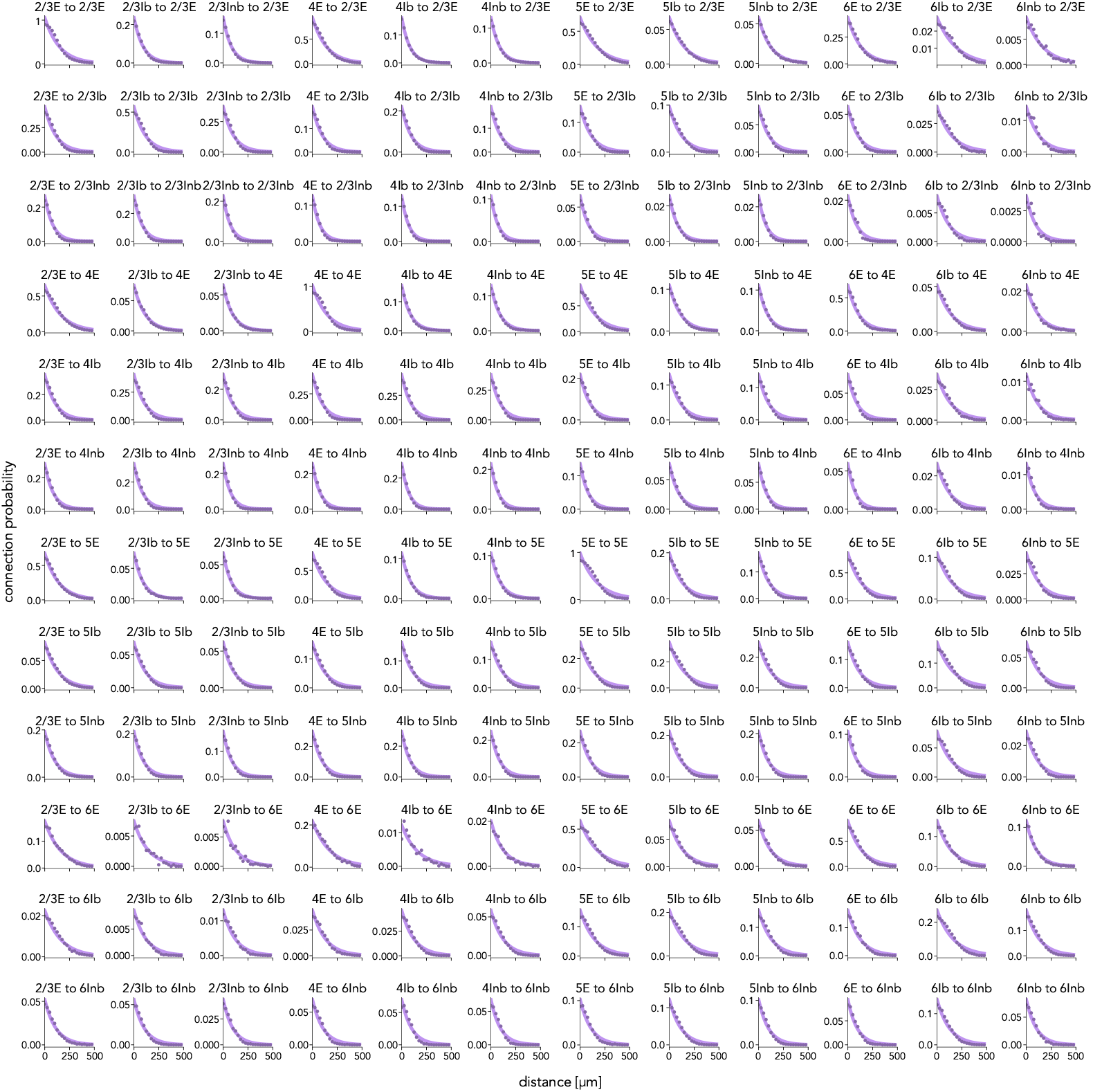
Connection probability between pre- and postsynaptic populations from EM data, potential connectivity. Purple dots indicate connection probability at distance *d*, purple curves are fits to the expected connection probability (Equation 2).

**Figure S4:**
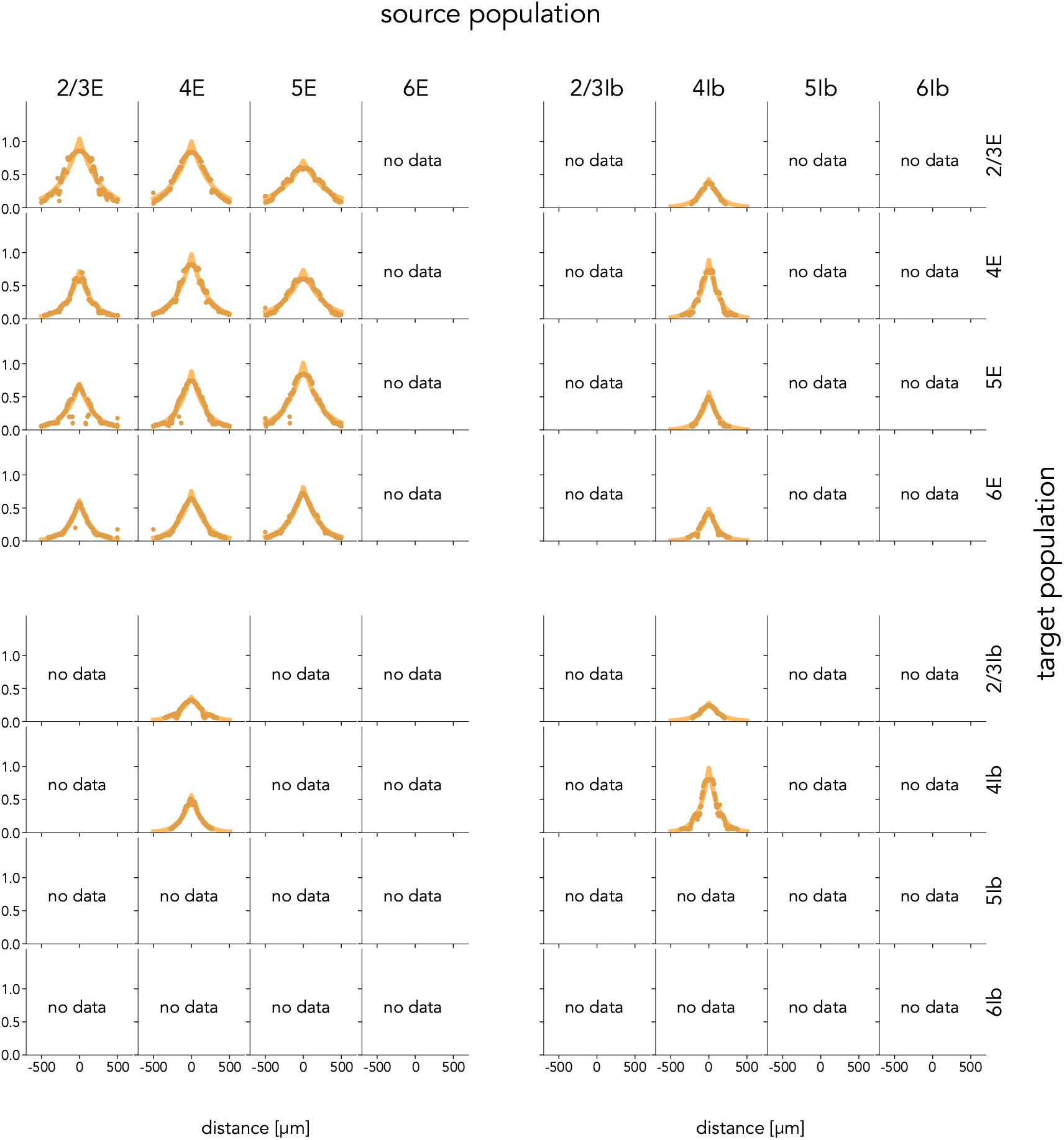
Potential connectivity between pre- and postsynaptic populations from LM data. Source population horizontal, target population vertical. Panels show potential connectivity (vertical) at a lateral displacement *d* as orange dots. Orange curves are fits to the connection probability according to Equation 2 for E-to-E connections, and mean number of synapses according to Equation 1 for all other types of connections.

**Figure S5:**
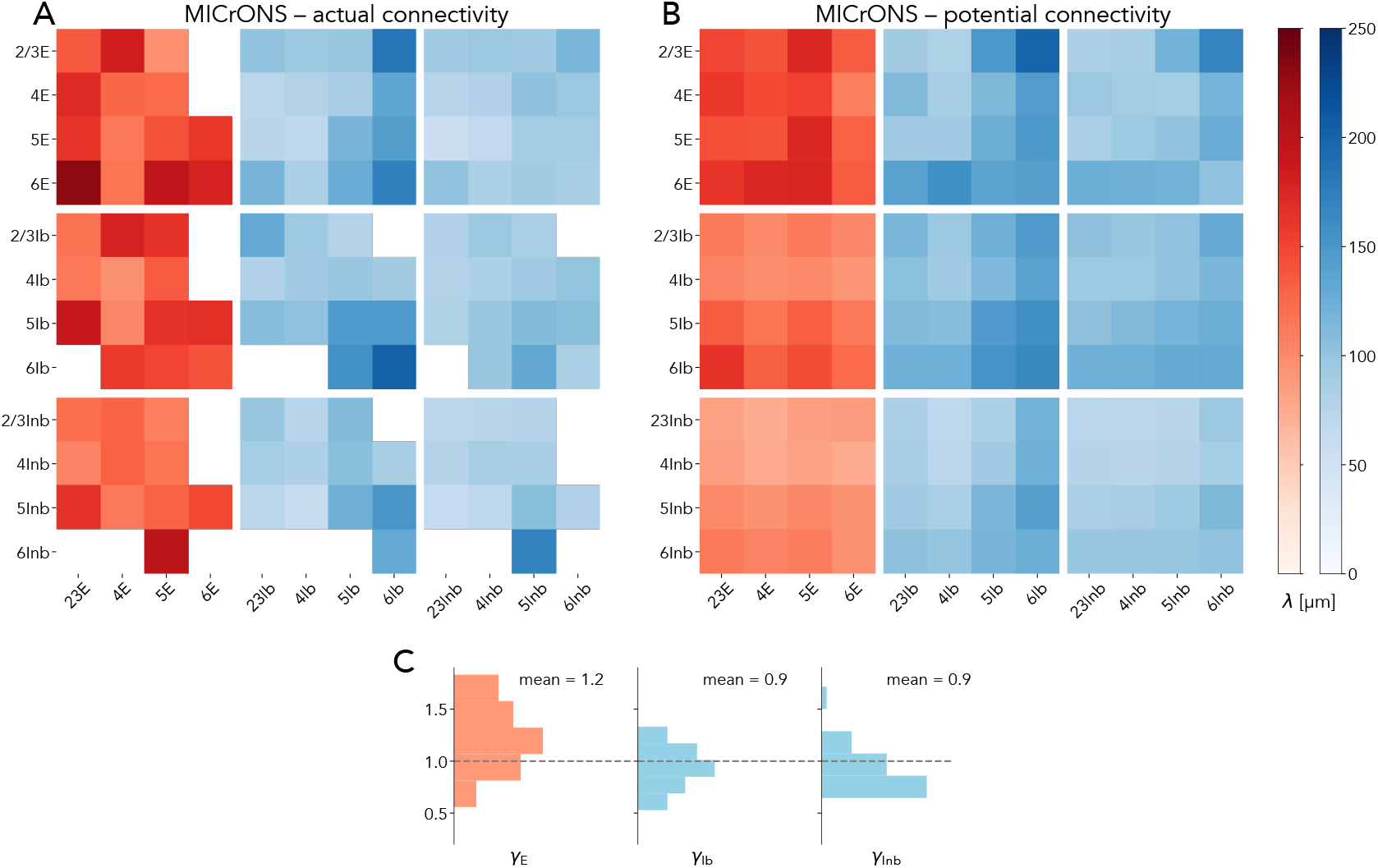
Comparison of the length scales of connection probability derived from actual and potential connectivity of EM data. **A** Characteristic length scale 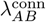 derived from actual connectivity. **B** Characteristiclength scale 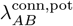 derived from potential connectivity. **C** Fraction of length scales 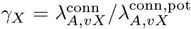 where *X* ∈ {E, Ib, Inb}.

### S1.3 Characteristic lengths for ℳ_LM_

Since the data of Stepanyants et al. [2] do not have the required resolution for the derivation in Section 2.2, we generalize the missing values from the estimated 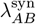. For inhibitory source populations, we choose

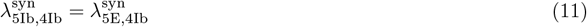

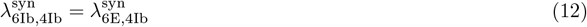

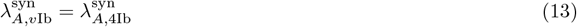

where *A* represents any target population, and *v* represents any source layer. Since Stepanyants et al. [2] only provide data for inhibitory basket cells, we use the same characteristic length scales also for inhibitory non-basket cells.

For excitatory source populations, we choose

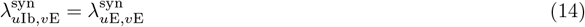

for all source layers except *v* = 6 since data for this layer is lacking. There, we choose

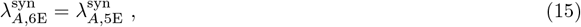

see Table 1.

**Table 1:**
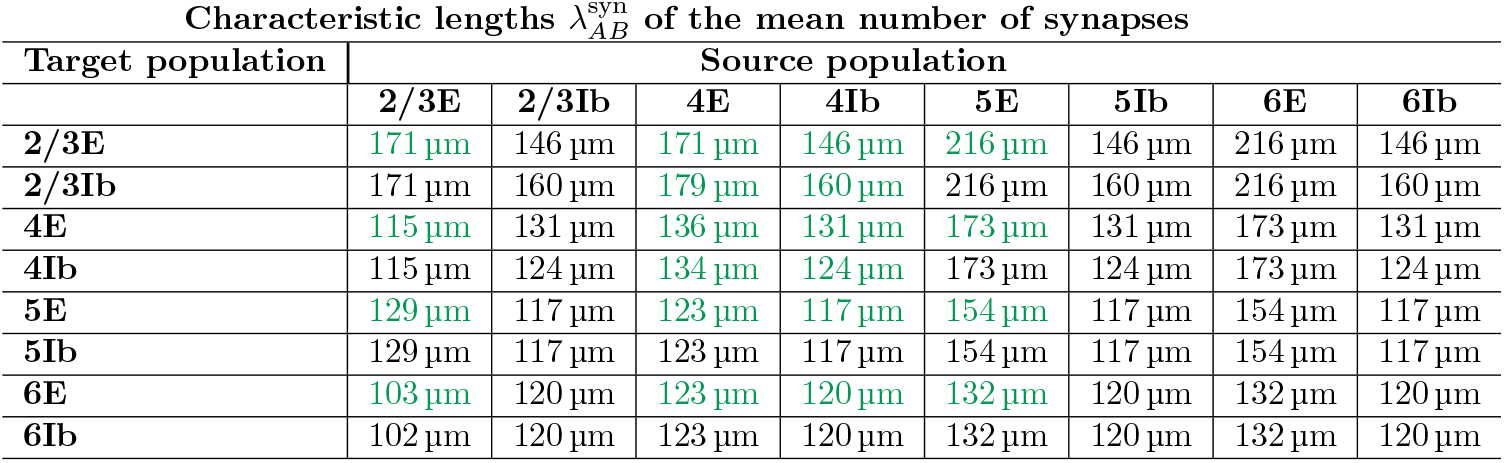
Characteristic lengths 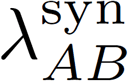 of the mean number of synapses for model ℳ_LM_. Green values are estimated directly from Stepanyants et al. [2], black values are generalized from these estimates.

### S1.4 Connection strengths

We follow Kraynyukova et al. [3] in defining the connection strength 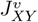 in a layer *v* from neurons of population *vY* to *vX* as

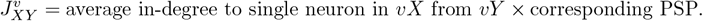

To compare our results with theirs, we only consider inhibitory basket cell populations and discard populations of inhibitory non-basket cells. We find that the weights chosen in the present work (c.f. Section 2.3) satisfy the hierarchy of connection strengths found in Kraynyukova et al. [3] (called connectivity weights in their study) in most cases. Additionally, we display the synaptic weights (assessed by unitary PSPs) used in Kraynyukova et al. [3] originating from previous measurements (their reference Allen Institute for Brain Science, 2019). For these synaptic weights, the hierarchy is satisfied for the EM model in all cases except L6 EI-EE (Figure S7, lower left panel). This is consistent with their findings using connection probabilities in mouse V1 obtained by electrophysiological means (their Figure 4).

**Figure S6:**
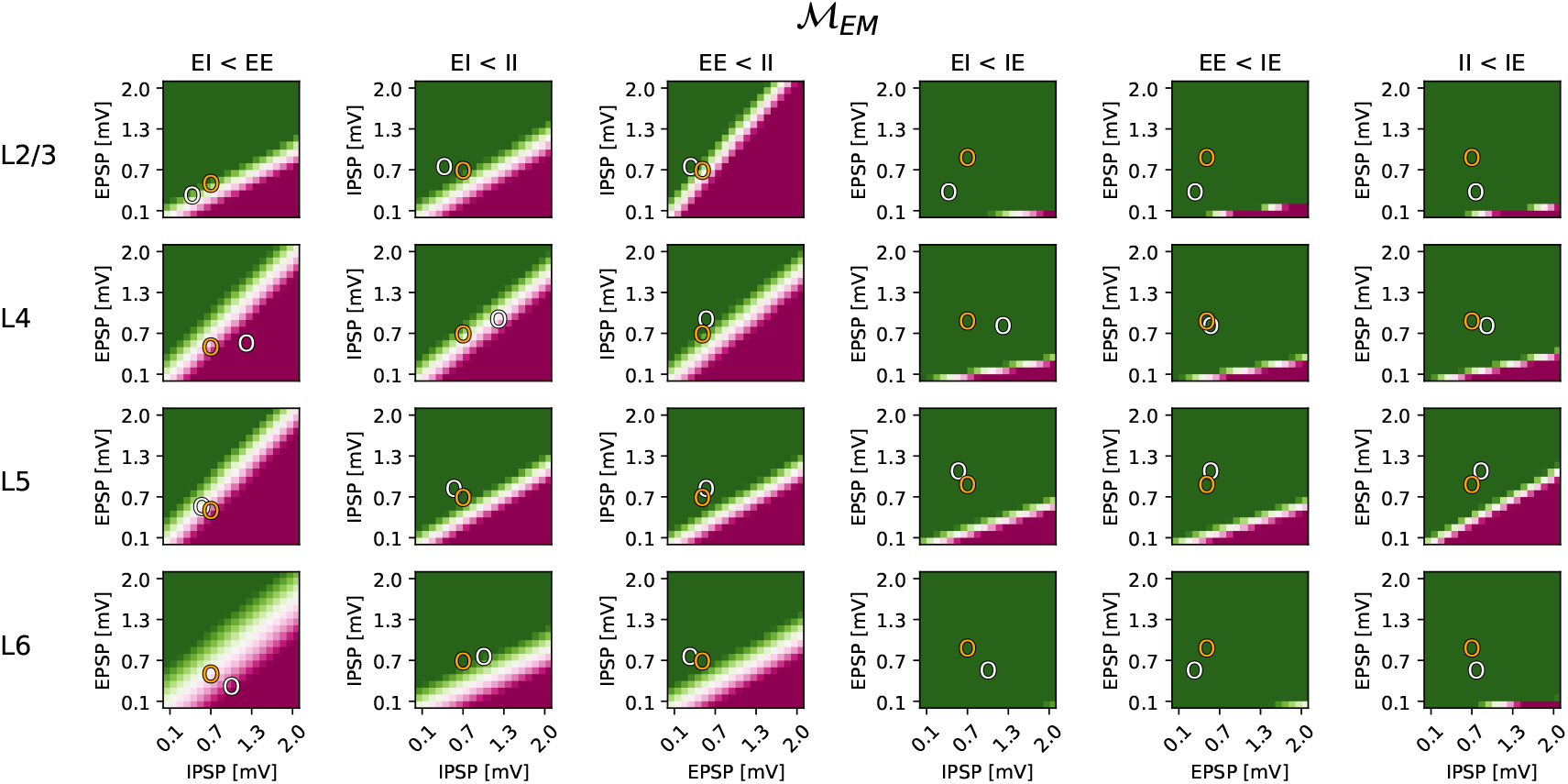
Connection strengths for ℳ_EM_. Layer- and population-pair-resolved connection strengths when systematically varying the PSP of presynaptic populations. On the horizontal (vertical) axis, the PSP of presynaptic populations with putatively smaller (larger) connection strengths is varied. Weights chosen in this work marked in orange, weights by the Allen Institute marked in white. Green (purple) indicates that the inequality given in the column title is fulfilled (unfulfilled) and therefore a hierarchy of connection strengths exists (does not exist).

**Figure S7:**
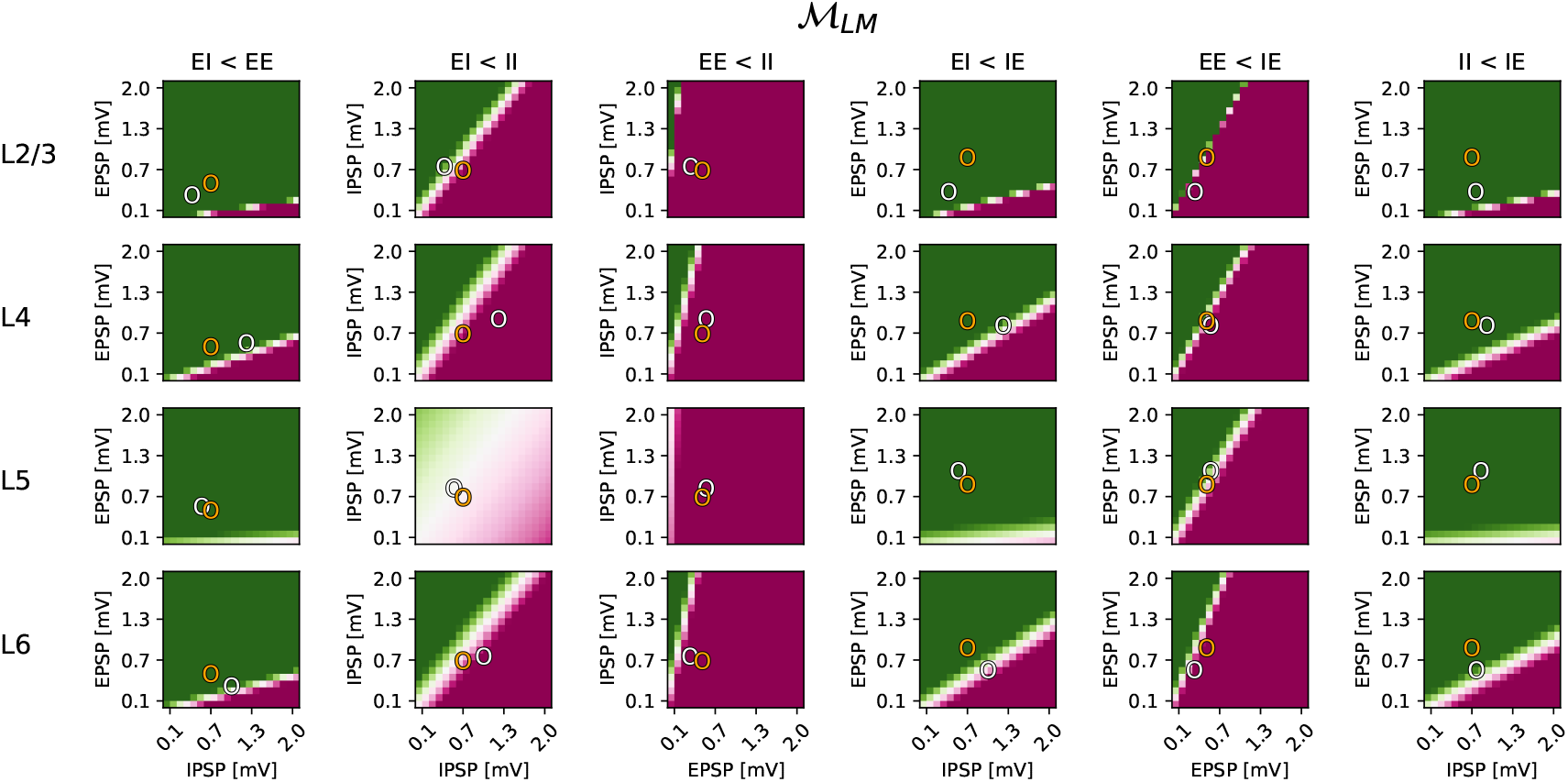
Connection strengths for ℳ_LM_. Same layout and color scheme as in Figure S6.

### S1.5 Tables summarizing model definitions

The network architecture and the list of parameters are summarized in the style of Nordlie et al. [4]. Table 2 summarizes the models, Table 3 specifies neuron and synapses models, Table 4, Table 5, and Table 6 give the numeric values used in the simulations.

**Table 2:**
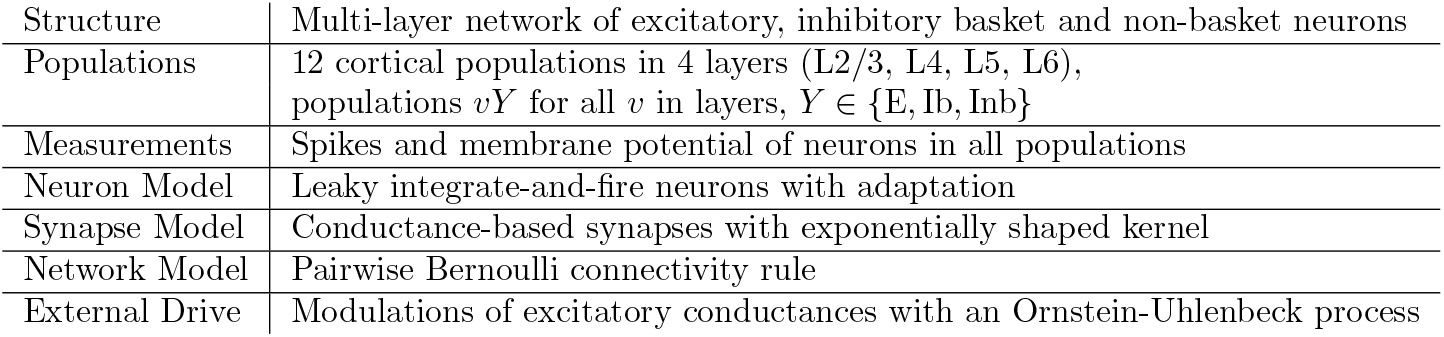
Model summaries for ℳ_LM_ and ℳ_EM_.

**Table 3:** Neuron and synapse models used for spiking neural network simulations of both ℳ_LM_ and ℳ_EM_.

| Neuron and synapse model |  |
| --- | --- |
| Sub-threshold dynamics | $C_m \dot{V}_m = -g_L (V_m - E_L) - (g_{ex} + g_{ext}) (V_m - E_{ex}) - g_{in} (V_m - E_{in}) - g_a (V_m - E_a)$ $\tau_{ex} \dot{g}_{ex}(t) = -g_{ex}(t) + \tau_{ex} \sum_j w_j \sum_{t_j} \delta(t - t_j - d_j)$ $\tau_{in} \dot{g}_{in}(t) = -g_{in}(t) + \tau_{in} \sum_k w_k \sum_{t_k} \delta(t - t_k - d_k)$ $\tau_a \dot{g}_a(t) = -g_a(t) + \tau_a q_a \sum_{t_i} \delta(t - t_i)$<br>where $t_i$ is the timing of the spike emitted by the neuron. |
| Spiking | If $V(t-) < V_{th}$ and $V(t+) \geq V_{th}$:<br>1. Set $t^* = t$ and $V(t) = V_{reset}$ in $(t^*, t^* + \tau_r]$.<br>2. Emit spike with time stamp $t^*$. |
| Computation time step |  |
| Time resolution | $dt = 0.1$ ms |
| Synaptic delays |  |
| Delay | Synaptic delay is uniformly distributed with a minimum of $dt$ and a maximum depending on the pre- and postsynaptic cell type. |
| Synaptic weights |  |
| Weights | Synaptic weights are lognormally distributed with a fixed relative (to the mean) standard deviation and the mean depending on the pre- and postsynaptic cell type. |
| Stimulation |  |
| External drive | $\tau_{ext} \dot{g}_{ext}(t) = -(g_{ext}(t) - \mu) + \sigma \sqrt{2\tau_{ext}} \eta(t)$<br>where $\eta$ denotes white noise, $\mu$ and $\sigma$ are population-specific. |

**Table 4:** Neuron parameters for both ℳ_LM_ and ℳ_EM_.

| Parameter | Value | Description |
| --- | --- | --- |
| **Shared neuron parameters** |  |  |
| $C_m$ | 250 pF | Membrane capacitance |
| $t_{\text{ref}}$ | 2 ms | Refractory period |
| $\tau_{\text{ex}}$ | 2 ms | Excitatory synaptic time constant |
| $\tau_{\text{in}}$ | 5 ms | Inhibitory synaptic time constant |
| $\tau_X$ | 10 ms | Time constant of Ornstein-Uhlenbeck noise |
| $g_L$ | 16.7 nS | Leak conductance |
| $E_L$ | −70 mV | Resting potential |
| $E_{\text{in}}$ | −75 mV | Inhibitory reversal potential |
| $E_{\text{ex}}$ | 0 mV | Excitatory reversal potential |
| $V_{\text{th}}$ | −50 mV | Threshold |
| $V_{\text{reset}}$ | −60 mV | Reset membrane potential |
| $V_m$ | −60 mV | Initial membrane potential |
| **Adaptation parameters: Excitatory neurons** |  |  |
| $q_a$ | 15.0 nS | Quantal spike-frequency adaptation conductance increase |
| $\tau_a$ | 110 ms | Time constant of spike-frequency adaptation |
| $E_a$ | −70 mV | Spike-frequency reversal potential |
| **Adaptation parameters: Inhibitory basket neurons** |  |  |
| $q_a$ | 4.0 nS | Quantal spike-frequency adaptation conductance increase |
| $\tau_a$ | 110 ms | Time constant of spike-frequency adaptation |
| $E_a$ | −70 mV | Spike-frequency reversal potential |
| **Adaptation parameters: Inhibitory non-basket neurons** |  |  |
| $q_a$ | 20.0 nS | Quantal spike-frequency adaptation conductance increase |
| $\tau_a$ | 110 ms | Time constant of spike-frequency adaptation |
| $E_a$ | −70 mV | Spike-frequency reversal potential |

**Table 5:** Maximum delays between populations of given cell types for both ℳ_LM_ and ℳ_EM_.

| **Maximum delay** |  |  |  |
| --- | --- | --- | --- |
| Target cell type | Source cell type |  |  |
|  | E | Ib | Inb |
| **E** | 2 ms | 0.8 ms | 1.5 ms |
| **Ib** | 1.2 ms | 1.5 ms | 1.5 ms |
| **Inb** | 1.5 ms | 1.2 ms | 1.5 ms |

**Table 6:** Mean connection weights between populations of given cell type for both ℳ_LM_ and ℳ_EM_. Numerical values inspired by measurements from Avermann et al. [5]. Relative standard deviation of weights equals 0.5.

| **Mean connection weights** |  |  |  |
| --- | --- | --- | --- |
| Target cell type | Source cell type |  |  |
|  | E | Ib | Inb |
| **E** | 1.0 nS | 11.5 nS | 9.5 nS |
| **Ib** | 2.0 nS | 11.5 nS | 7.5 nS |
| **Inb** | 1.0 nS | 15.5 nS | 9.5 nS |

**Table 7:** Mean of conductance fluctuations modeled as OU processes providing external drive for the tuned ℳ_LM_ network.

| $\mu_X$ for $\mathcal{M}_{\text{LM}}$ | | | |
| --- | --- | --- | --- |
| Layer | Cell type |  |  |
|  | E | Ib | Inb |
| **L2/3** | 3.90 nS | 3.82 nS | 5.02 nS |
| **L4** | 4.78 nS | 3.91 nS | 5.31 nS |
| **L5** | 4.68 nS | 4.13 nS | 4.72 nS |
| **L6** | 4.22 nS | 3.98 nS | 4.95 nS |

**Table 8:** Mean of conductance fluctuations modeled as OU processes providing external drive for the tuned ℳ_EM_ network.

| $\mu_X$ for $\mathcal{M}_{\text{LM}}$ | | | |
| --- | --- | --- | --- |
| Layer | Cell type |  |  |
|  | E | Ib | Inb |
| **L2/3** | 7.34 nS | 4.48 nS | 5.66 nS |
| **L4** | 8.51 nS | 7.25 nS | 7.39 nS |
| **L5** | 12.52 nS | 14.86 nS | 9.39 nS |
| **L6** | 5.26 nS | 3.24 nS | 4.34 nS |

